## Supplementary figures and images for "Zebrafish macrophage developmental arrest underlies depletion of microglia and reveals Csf1r-independent metaphocytes"

### Supplemental Figure 1

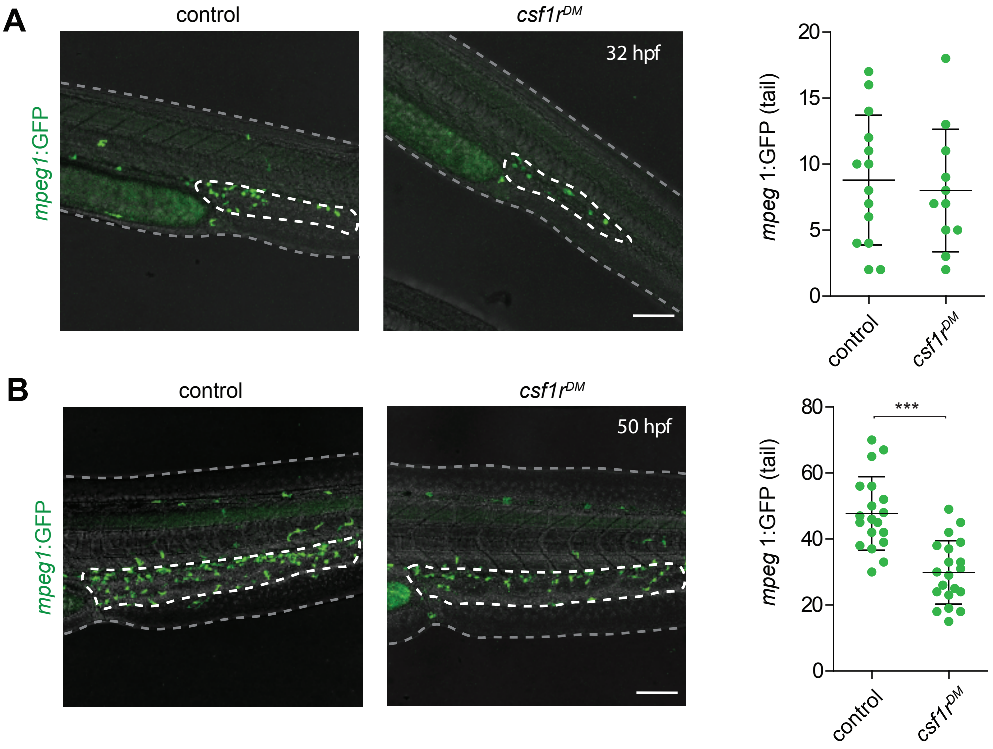

### Supplemental Figure 2

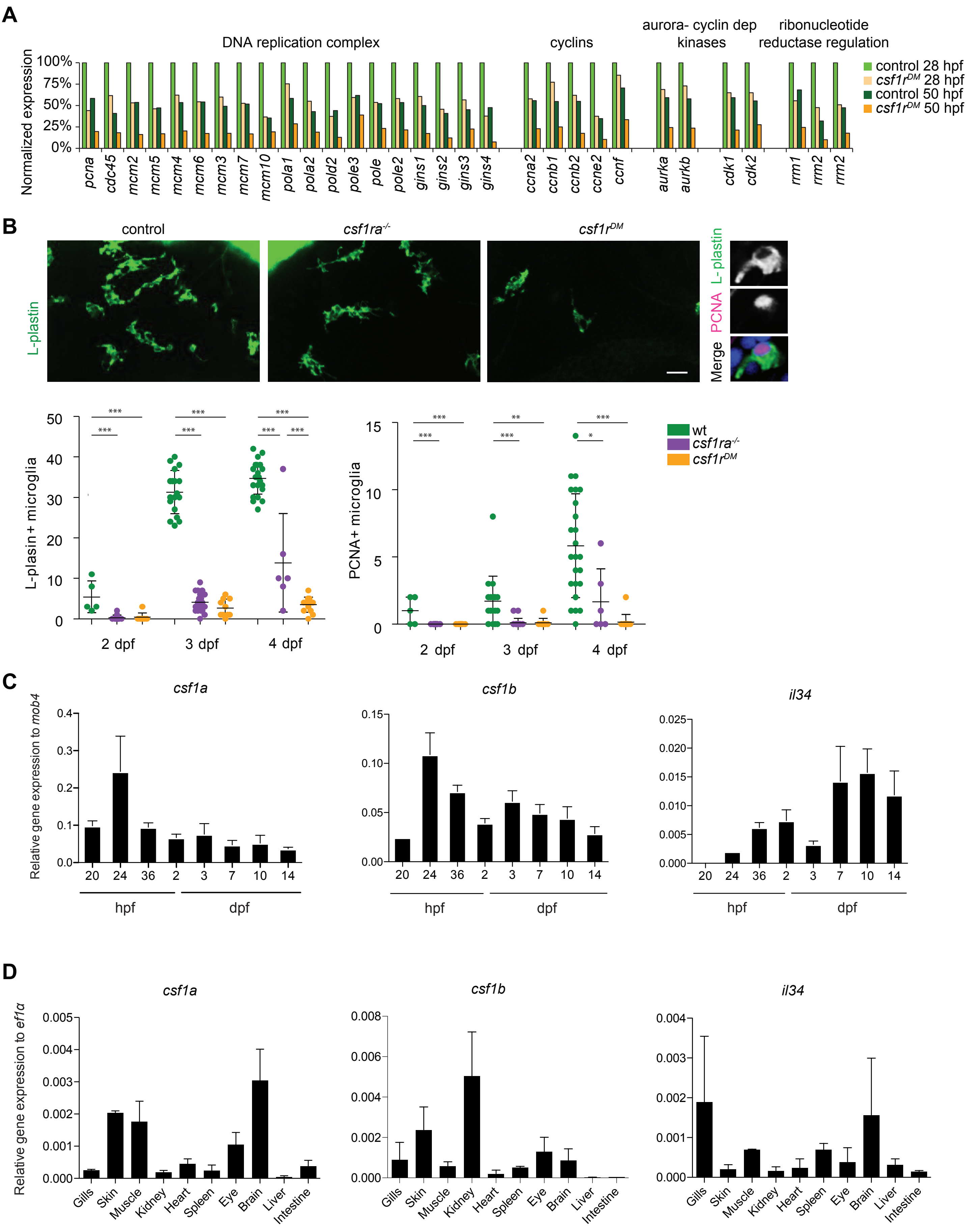

### Supplemental Figure 3

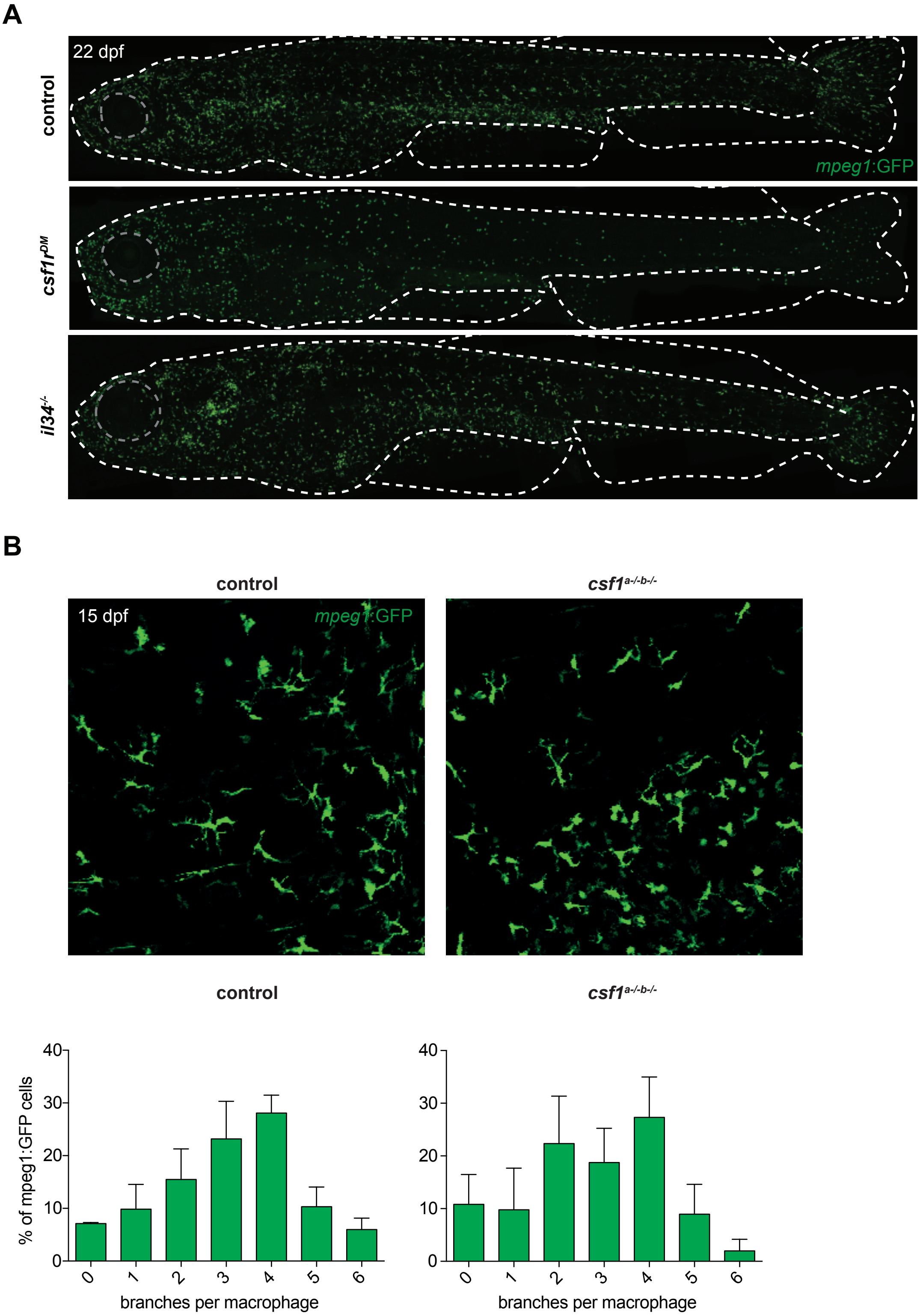

### Supplemental Figure 4

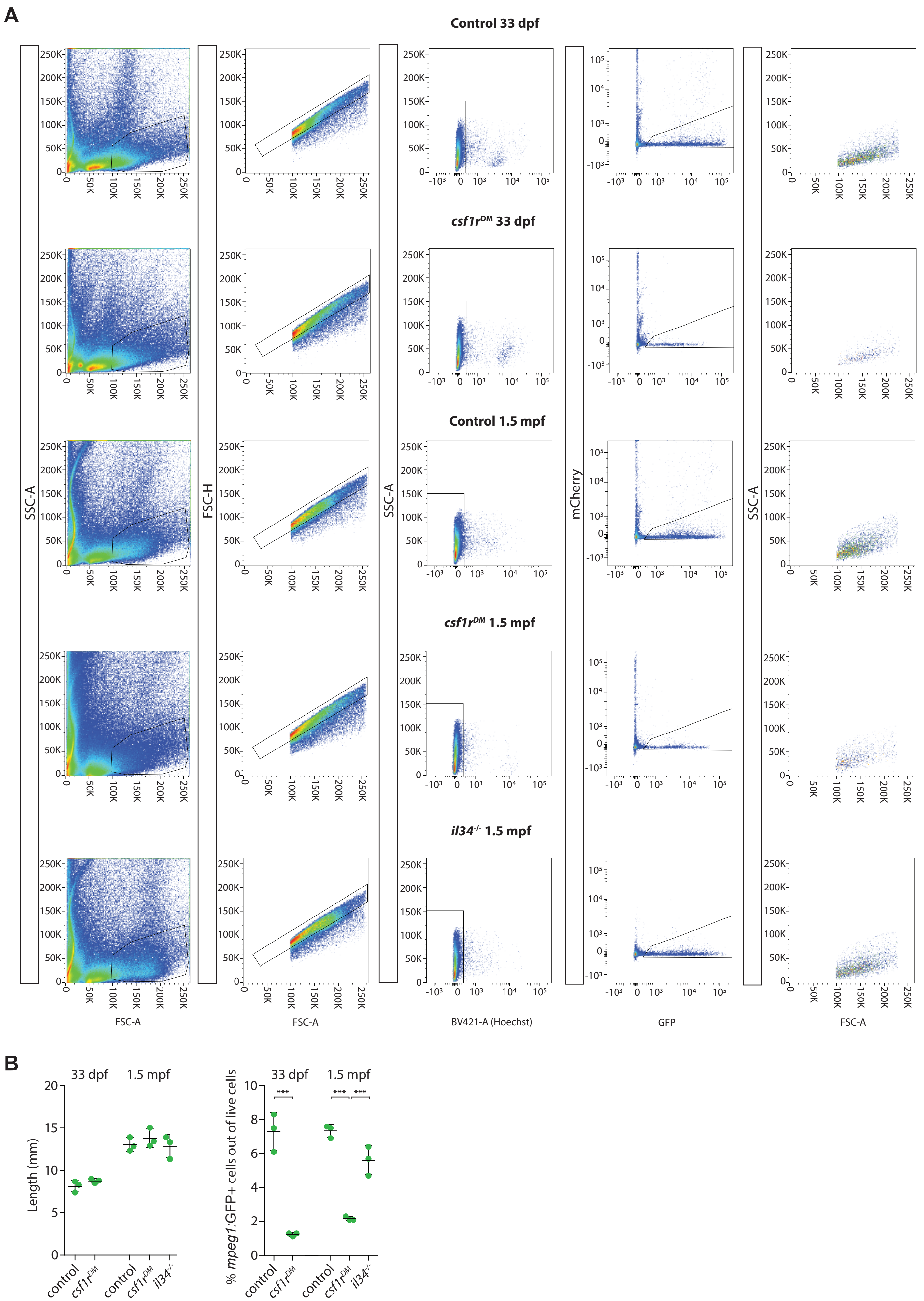

### Supplemental Figure 5

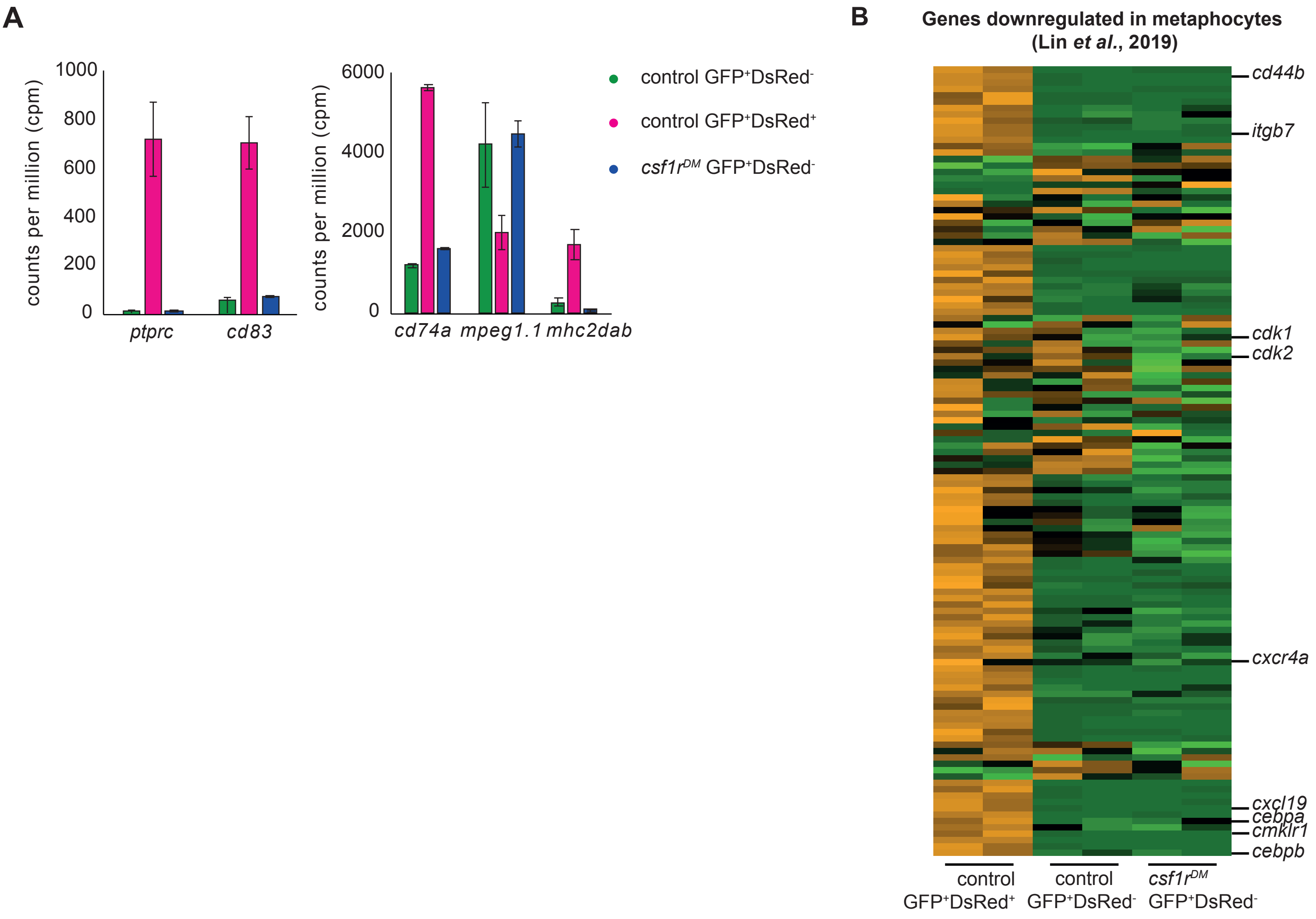
